# Evaluation of Fendiline Treatment in VP40 System with Nucleation-Elongation Process: A Computational Model of Ebola Virus Matrix Protein Assembly

**DOI:** 10.1101/2023.08.03.551833

**Authors:** Xiao Liu, Monica Husby, Robert V. Stahelin, Elsje Pienaar

## Abstract

Ebola virus (EBOV) infection is threatening human health, especially in Central and West Africa. Limited clinical trials and the requirement of biosafety level-4 (BSL-4) laboratories hinders experimental work to advance our understanding of EBOV and evaluation of treatment. In this work, we use a computational model to study the assembly and budding process of EBOV and evaluate the effect of fendiline on these processes. Our results indicate that the assembly of VP40 filaments may follow the nucleation-elongation theory, as it is critical to maintain a pool of VP40 dimer for the maturation and production of virus-like particles (VLPs). We further find that the nucleation-elongation process can also be influenced by phosphatidylserine (PS), which can complicate the efficacy of fendiline, a drug that lowers cellular PS levels. We observe that fendiline may increase VLP production at earlier time points (24 h) and under low concentrations (≤ 2 μM). But this effect is transient and does not change the conclusion that fendiline generally decreases VLP production. We also conclude that fendiline can be more efficient at the stage of VLP budding relative to earlier phases. Combination therapy with a VLP budding step-targeted drug may further increase the treatment efficiency of fendiline. Finally, we also show that fendiline has higher efficacy when VP40 expression is high. While these are single-cell level results based on the VP40 system, it points out a potential way of fendiline application affecting EBOV assembly, which can be further tested in experimental studies with multiple EBOV proteins or live virus.

**Importance:** EBOV infection can cause deadly hemorrhagic fever, which has a mortality rate around 90% without treatment. The recent outbreaks in Uganda and Democratic Republic of the Congo illustrate its treat to human health. Though two antibody-based treatments are approved, mortality rates in the last outbreak is still higher than 30%. This can partly be due to the requirement of advanced medical facilities for current treatments. As a result, it is very important to develop and evaluate new therapies for EBOV infection, especially those can be easily applied in the developing world. The significance of our research is that we evaluate the potential treatment effect of fendiline on EBOV infection in the VP40 system with a computational approach, which both greatly saves time and lowers cost compared to traditional experimental studies, and provides innovative new tools to study viral protein dynamics.

## Introduction

Ebola virus (EBOV) causes hemorrhagic fever, a fatal disease with a high mortality rate in humans (1, 2). Since the discovery of EBOV in 1976, it has caused more than 34,000 cases and 15,000 deaths (3). New treatments of EBOV are being developed, and two monoclonal antibody therapies (Inmazeb and Ebanga) were approved by the FDA in 2020. However, even with treatment, the mortality rate is still higher than 30%, and side effects can be severe (4–6). Moreover, monoclonal antibodies, as a large protein molecule, need to be applied through intravenous (IV) infusion or injections. Advanced medical facilities and equipment required for such therapies may not be available in affected areas. In 2022, an estimated one third of infected patients died from EBOV disease (3). Thus, there is a significant need to develop more effective and accessible treatments for Ebola virus disease (EVD).

Fendiline is a calcium channel blocker used for arrhythmic and anginal diseases (7, 8) and has recently been proposed as a potential anti-viral therapy for EBOV (9). Studies have shown that fendiline reduces the PS content in the cellular plasma membrane inner leaflet by inhibiting the activity of acid sphingomyelinase (ASM) (10, 11). Plasma membrane PS levels are critical for the production of EBOV VP40 virus-like particles (VLPs), as it will influence VP40 dimer membrane association and oligomerization (12, 13). Our previous work also suggested that the VLP budding step (process of mature VLP detaching from the cell surface) could be influenced by PS (14). As a result, fendiline could reduce EBOV VP40 VLP production (15), thus significantly reducing EBOV replication. However, detailed mechanistic results from experimental fendiline treatment are still lacking.

A cellular system using VP40-based VLPs is a valuable system for studying the assembly and budding process of EBOV. VP40, the matrix protein of EBOV, can assemble into filaments and form VLPs when expressed in the absence of other EBOV proteins in mammalian cells (12, 16–18). However, our knowledge of the mechanistic aspects of the assembly process is still emerging. For example, the VP40 dimer was identified as the building block for VP40 oligomers/filaments at different plasma membrane assembly sites (19). We also lack understanding of VP40 assembly and VLP budding dynamics, as well as the regulation mechanism behind the VLP production process. For example, the nucleation-elongation theory has been proposed for other filamentous oligomerization processes (20–22), such as amyloid fibers (23), actin (24), myosin (25) and DNA nanotubes (26). Our previous computational work suggested that the same nucleation-elongation process could be at play in VP40 filament growth (14).

Computational studies have a long history of complementing experimental studies in biological and medical areas, as in-silico or “virtual experiments” can be conducted in quicker time frames and integrate information from diverse data sources. It is not new to evaluate small molecule treatment of diseases (27–29) or to study the nucleation-elongation theory (30) by computational methods. However, neither of these principles has been applied to EBOV at the intracellular level. Thus, we aim to take advantage of computational approaches to explore if the nucleation-elongation process applies in VP40 filament growth. We also test and evaluate the ability of fendiline to impact VLP production with this built-in nucleation-elongation mechanism. By doing so, we will complement experimental studies with computational approaches.

In this study, we incorporated the nucleation-elongation process for VP40 filament oligomerization into our existing ordinary differential equation-based (ODE-based) model of EBOV VP40 assembly and budding (14). We then applied our model to simulate fendiline treatment, and evaluated the impact of different fendiline concentrations, application timing and co-treatment with other hypothesized treatments on VP40 VLP production. Our simulations provided quantitative insights into the dynamics of VP40 assembly and VLP production, as well as the impact of fendiline on these processes.

## Results

### Nucleation-elongation assembly, and direct PS influence on this process, are required to reproduce observed relative oligomer frequency in the VP40 VLP system

Our previous work showed that without a mechanism resembling nucleation-elongation, the decreasing relative VP40 oligomer frequency with oligomer size and VLP production data cannot be simultaneously reproduced by the model (14). Thus, we proposed that the nucleation-elongation process exists in VP40 filament assembly, similar to other oligomers (14, 23–26). To explore the implications of nucleation-elongation for VP40 assembly dynamics, we explicitly incorporated the nucleation-elongation process into our existing model, which replaced the “filament stabilization” process in the original model (14). We named this model the ‘As0’ model and calibrated the model to published experimental data as described in the methods section “Parameter estimation and calibration” as well as prior work (14). The ‘As0’ model successfully reproduces experimental data measuring oligomer ratio, VLP production, VP40 budding ratio, VP40 plasma membrane localization, relative VLP production (Fig. 1A-E) as well as the decreasing trend of relative oligomer frequency from cell membrane VP40 dimer to 42mer (Fig. 1F).

**Figure 1.**
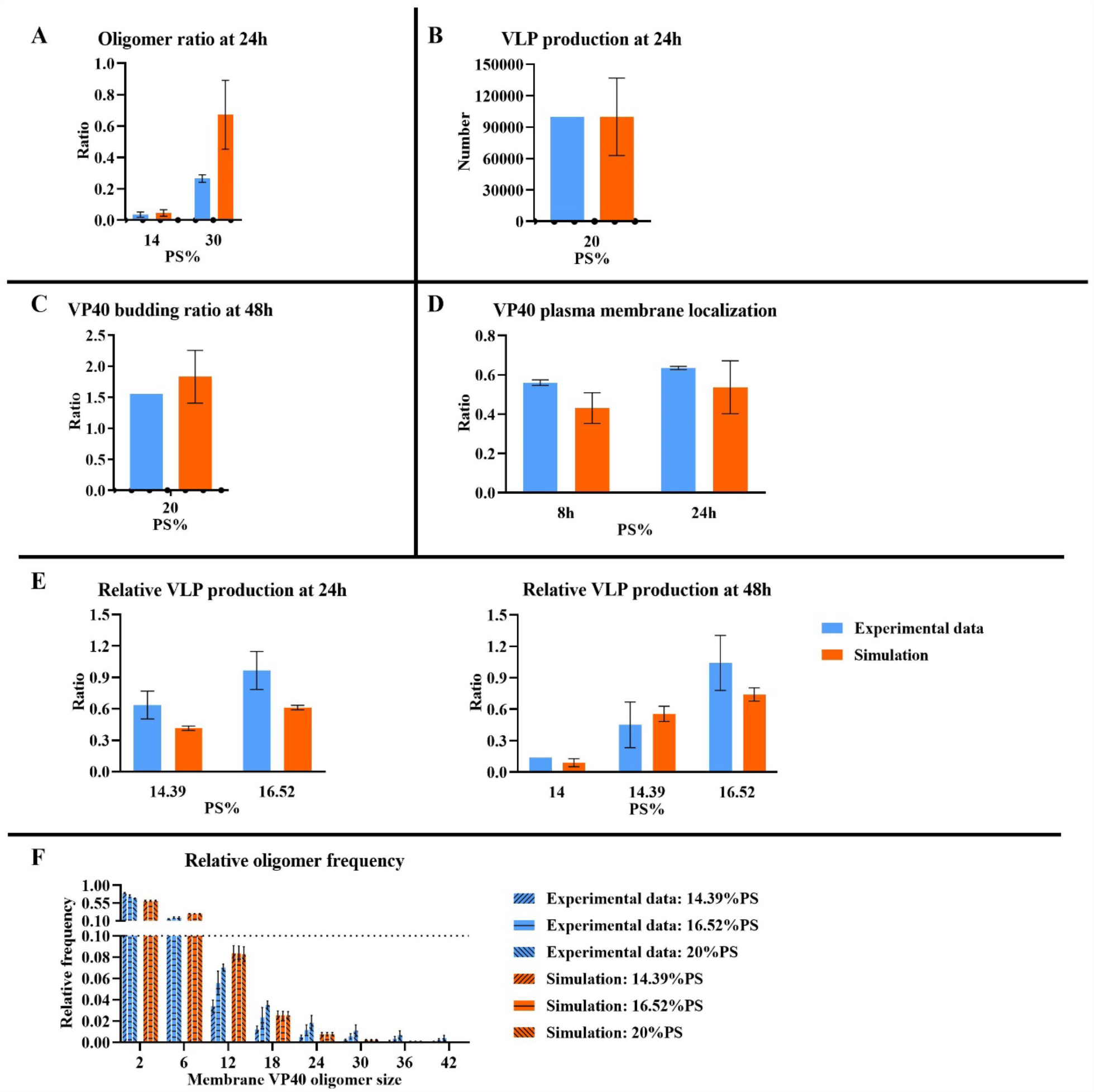
Simulation result from the ‘As0’ model. (A) Oligomer ratio at 24h. (B) VLP production at 24h. (C) VP40 budding ratio at 48h. (D) VP40 plasma membrane localization. (E) Relative VLP production. (F) Relative oligomer frequency. While the deceasing trend of relative frequency from membrane VP40 dimer to 42mer is predicted, the increasing trend in higher oligomers under higher PS level is not reproduced. The three bars in each of the sub-column are 14.39%, 16,52%, 20% PS from left to right separately. Error bars indicate SEM from top 5 fits.

However, in the absence of any direct PS influence on the nucleation elongation mechanism, the increase in oligomer frequency with increasing PS levels is not reproduced for larger oligomers (Fig. 1F). Similar to our prior approach (14), we addressed this limitation of model ‘As0’ by assuming a direct influence of PS on the nucleation-elongation process. To characterize the nature of PS influence on nucleation and/or elongation steps, we test both positive and negative influence of PS on nucleation or elongation processes in models ‘As1’-’As4’ (Table 1, Eq. 15-16). We evaluated the ability of each model to reproduce the experimentally observed trend that higher PS levels results in increased relative frequency of larger oligomers. Predictions from models ‘As1’ and ‘As4’ show the opposite trend in higher oligomers (6-42mers) compared to experimental data (Fig. 2A, D), with a slight decrease in frequency of larger oligomers as PS increases. Predictions from ‘As2’ and ‘As3’ match the trend observed in experimental data (Fig. 2B-C). Both ‘As2’ and ‘As3’ can still reproduce the other experimental data sets, and therefore did not lose accuracy compared to ‘As0’ (Fig. S1, S2).

**Table 1.**
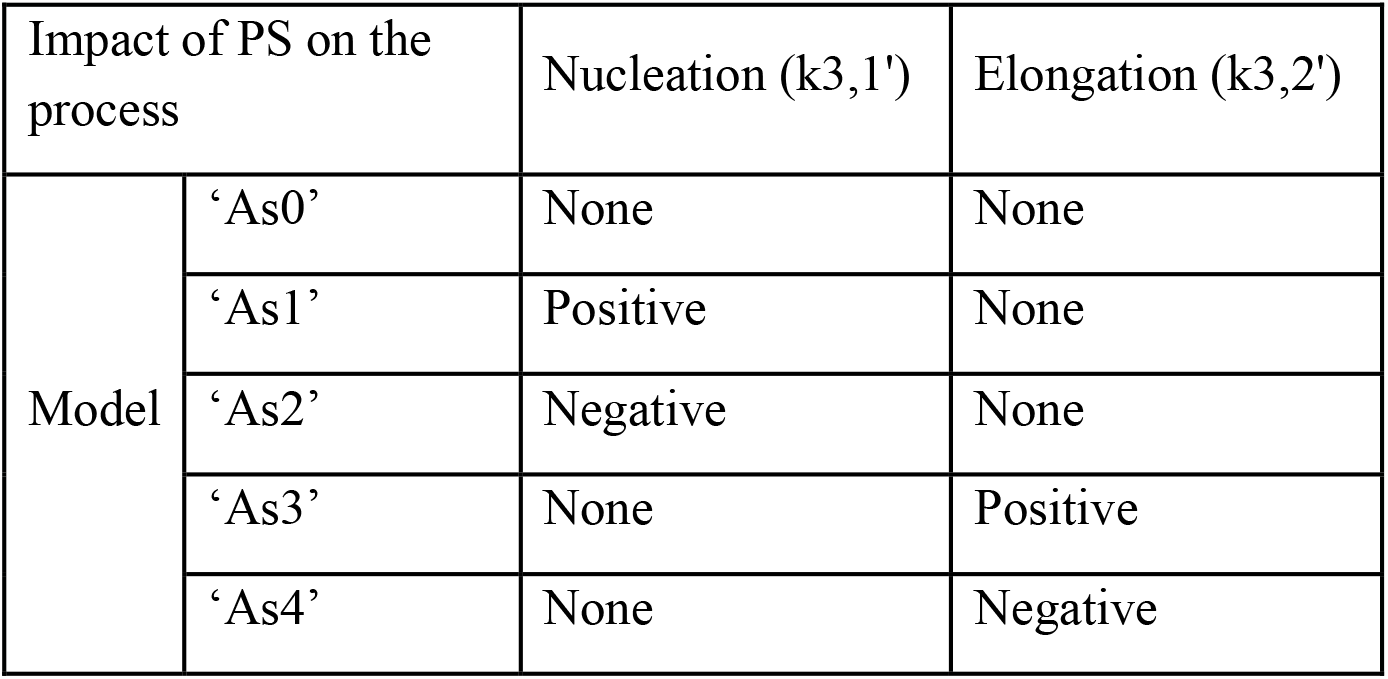
Model construction.

**Figure 2.**
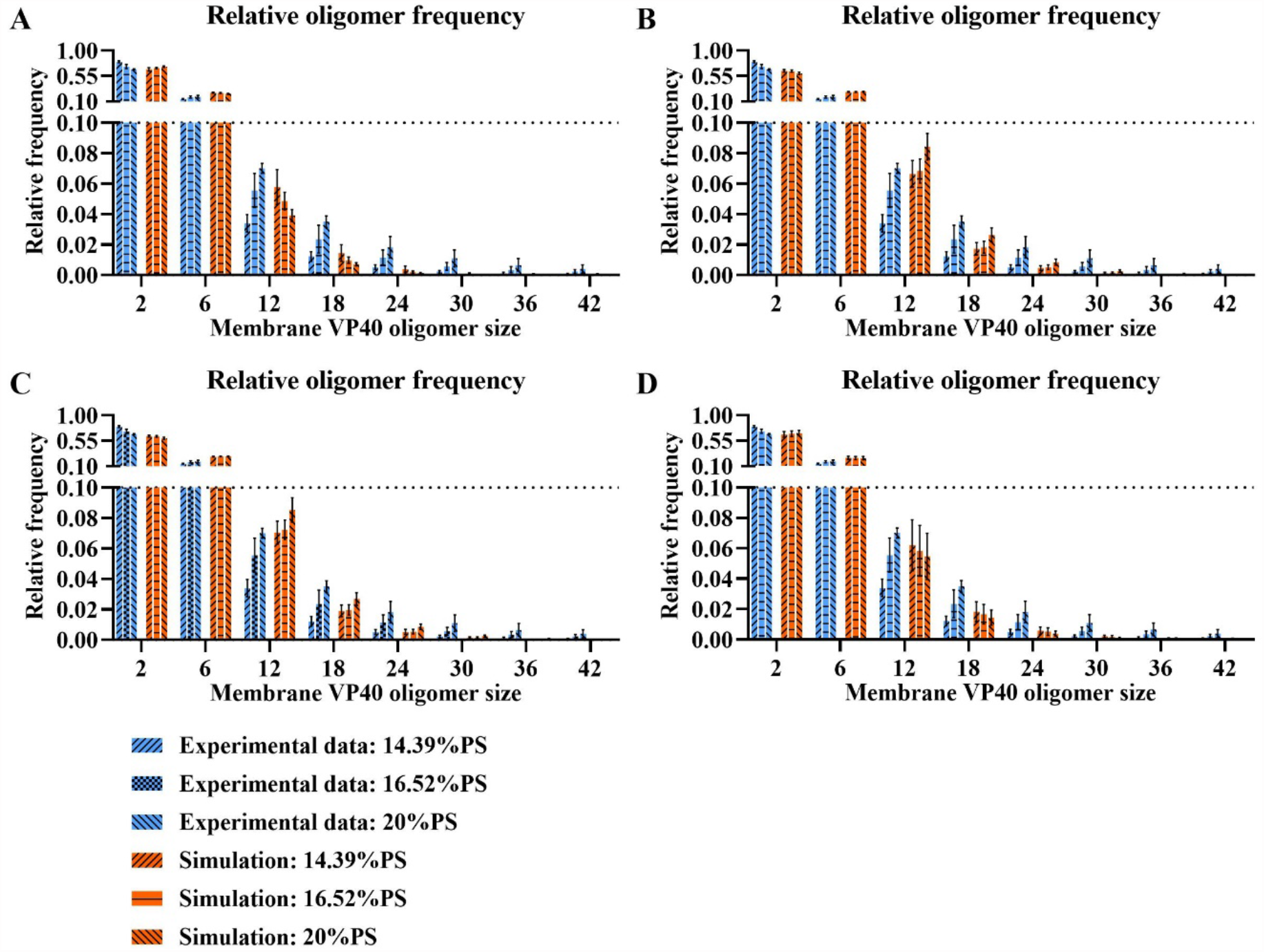
Simulation of relative oligomer frequency for ‘As1’-’As4’ models. (A) ‘As1’ model fails to predict the trend. (B) ‘As2’ model predicts the trend successfully. (C) ‘As3’ model predicts the trend successfully. (D) ‘As4’ model fails to predict the trend. Error bars indicate SEM from top 5 fits.

While ‘As2’ and ‘As3’ models have different PS influence on the nucleation-elongation process (Table 1), there is a common feature between the two models. The reverse rate constant of elongation (k_3,2_’) is by definition always lower than the reverse rate constant of nucleation (k_3,1_’), representing the stabilization of growing oligomers. Thus, a decrease in the reverse rate constant for nucleation (as in ‘As2’) or an increase in the reverse rate constant for elongation (as in ‘As3’) will both decrease the difference between nucleation and elongation processes under higher PS levels. This result suggests that the increase in relative frequency of larger oligomers is related to the decreased difference between nucleation and elongation reverse rate constants (k_3,1_’ and k_3,2_’) as PS levels increase. Taken together, our findings indicate that VP40 assembly through a nucleation-elongation mechanism is consistent with experimental measurements; and that high PS levels diminish the difference between the nucleation and elongation phases resulting in higher frequencies of larger oligomers.

### Fendiline treatment simulation detects rare cases where fendiline increases VLP production

Having confirmed that our model can reproduce the influence of PS on VP40 VLP assembly and budding, we next aimed to simulate fendiline treatment and evaluate its effects on VP40 VLPs in the context of the nucleation-elongation dynamics. To produce a distribution of biologically feasible simulations and account for parameter uncertainty, we performed LHS sampling within the parameter ranges identified during model calibration. We selected all (75 out of 950 in ‘As2’ and 50 out of 950 in ‘As3’ model) parameter sets with a cost lower than 3 or score higher than 5 for fendiline treatment simulation as described in the methods section “Parameter estimation and calibration”.

Our first simulation applied 0.5-10 μM of fendiline to the chosen parameters and models. Our simulations show that in most cases, as fendiline concentration increases, VLP production decreases as a result of fendiline-driven reduction in plasma membrane PS levels (Fig. 3, Table S1-S2). This is consistent with our expectation (based on model structure) and experimental observations (13, 15).

**Figure 3.**
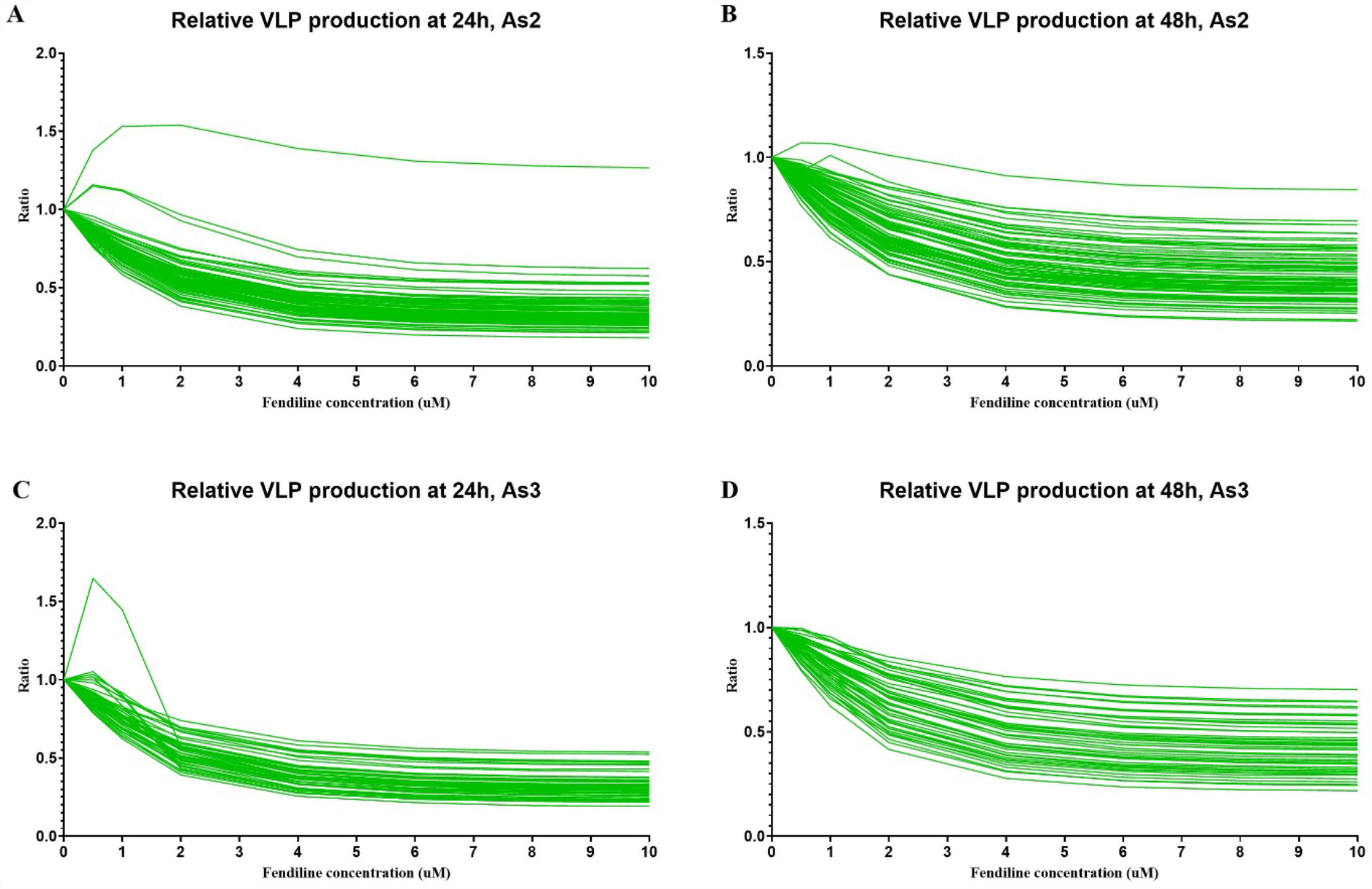
Relative VLP production in fendiline simulation for ‘As2’ and ‘As3’ models. (A) Relative VLP production at 24h for ‘As2’ model. (B) Relative VLP production at 48h for ‘As2’ model. (C) Relative VLP production at 24h for ‘As3’ model. (D) Relative VLP production at 48h for ‘As3’ model.

However, when the fendiline concentration is low (< 2 μM), several parameter sets in both ‘As2’ and ‘As3’ show an increase in VLP production over the short-term (24h). The elevation in VLP becomes less pronounced or reversed when fendiline concentration increases beyond 2 µM (Fig. 3A, C). For the longer-term (48 h), the fendiline induced increase in VLP production becomes less pronounced (Fig. 3B, D).

Thus, our simulations indicate that while fendiline generally lowers VLP production, it has the potential to counterintuitively increase VLP production at concentrations < 2 μM. This finding is consistent with relative VLP production experimental data (15).

### Fendiline can increase VLP production under slow filament growth and high VLP budding rate conditions

To identify potential mechanisms behind the counterintuitive ability of fendiline to increase VLP production, we enriched our parameter sampling around the parameter set in ‘As2’ which shows increased VLP production at 24 h and 48 h under 2 μM of fendiline treatment (Fig. 3A, B). Since the ‘As2’ and ‘As3’ models appear to behave similarly in fendiline simulations, and the ‘As2’ model has more samples that capture a broader diversity of the biologically observed dynamics (elevated VLP production at higher concentration of fendiline), our further simulations will only use the ‘As2’ model. We sampled 1000 new parameter sets around the parameter set of interest (Table S3). Parameter sets that have no VLP production at the time of evaluation in the absence of fendiline treatment were excluded from the analysis. Our results show that 199 out of 656 fendiline-treated simulations have elevated VLP production at 24 h, and 147 out of 955 have elevated VLP production at 48 h under 2 µM fendiline (Table S4).

When analyzing the parameter distributions of the simulations in which 2 μM fendiline causes an increase in VLP production at 48 h, we identified three important parameter conditions: low k_3_ (filament growth forward rate constant), low k_D3,1_ (nucleation equilibrium constant) as well as high k_4_ (VLP budding rate constant) (Fig. 4, Table S5). Considering these findings in the context of our model structure, we hypothesize that a low k_3_ and k_D3,1_, which indicates an even lower k_3,1_’, will slow down the maturation of filaments and postpone the starting time of VLP production. The impact of k_3,1_’ may be counterintuitive. But the mechanism behind this is that it allows more membrane dimers to accumulate in small-sized filaments and thus decreases the building block for large filament growth, which is similar to the observation from our previous work (14) as well as studies demonstrating VP40 assembly occurring at different patches in the plasma membrane (16, 19, 31). However, the application of fendiline will decrease PS levels, thus elevating k_3,1_’ (As the mechanism of ‘As2’ model) and resulting in earlier VLP production compared to untreated cases. Moreover, a high k_4_ means that the budding step is not rate-limiting, and therefore the fendiline-induced reduction in PS has minimal impact on the VLP budding step.

**Figure 4.**
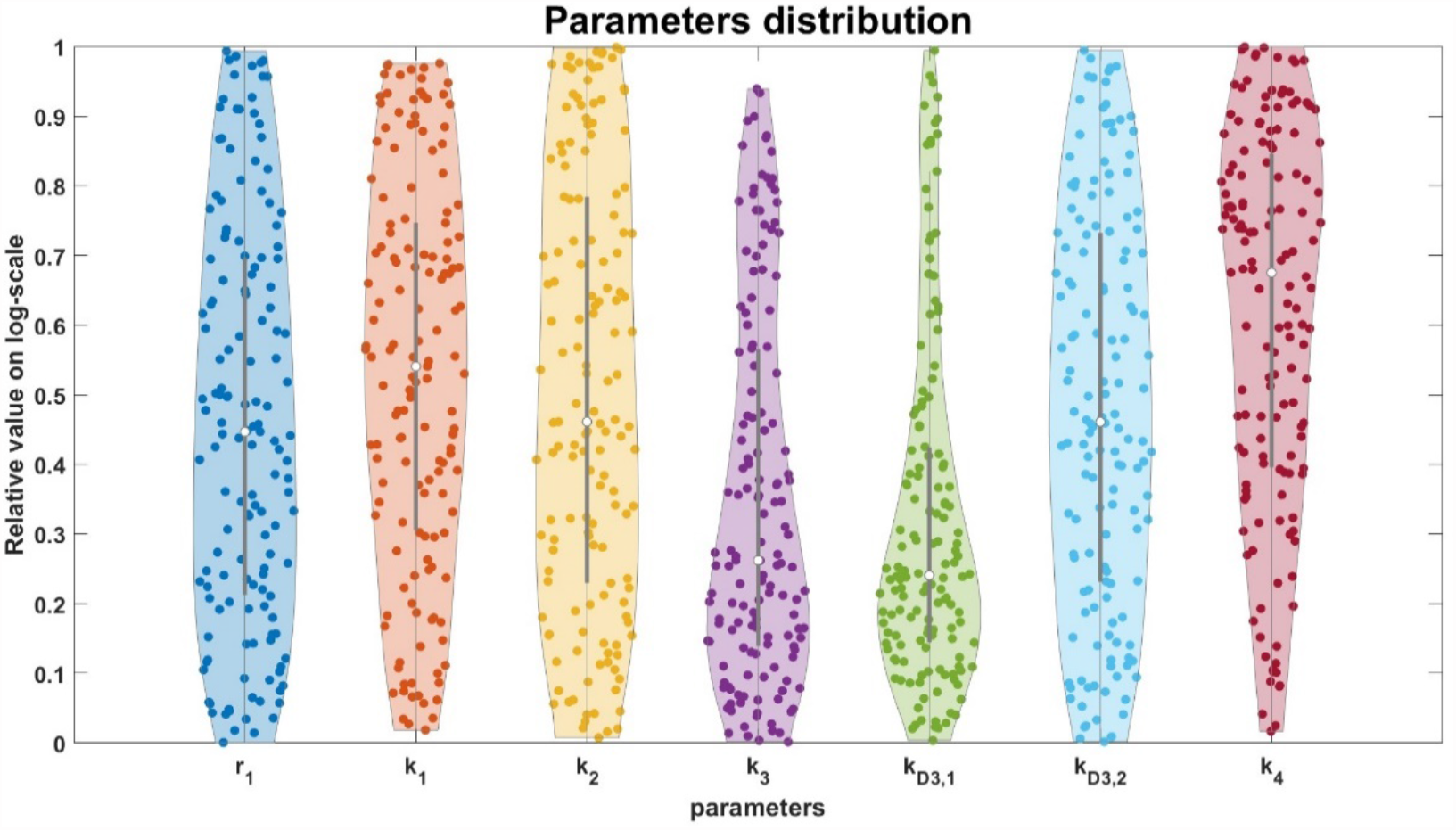
Parameter distributions where fendiline results in increased VLP production. Low k_3_, low k_D3,1_, high k_1_ and high k_4_ are related to fendiline induced VLP increase. The Y axis range shows the relative value of each parameter in their LHS. 0 indicates lower bound and 1 indicates upper bound.

To confirm our hypothesis on the impact of these parameters and further explore mechanisms behind it, we then looked at the individual dynamics of the system. We divided all 955 “effective” parameter sets into 2 types: a late VLP production type where VLPs start being produced at 40 h or later, and an early VLP production type where VLPs start being produced prior to 40 h. In late VLP production type simulations, most parameter sets (73 out of 75) show increases in VLP production under fendiline treatment, which is due to fendiline driving earlier VLP production (Fig. 5A). These parameter distributions are characterized by very low k_3_ and k_D3,1_ (Fig. S3A, Table S5). In early VLP production type, only a small portion of the simulations (74 out of 880) has increased VLP production under 2 μM fendiline treatment. The increased VLP production in this type is mostly caused by fluctuations. Fendiline therefore, cannot compensate for fluctuations in VLP production (Fig 5B). This is also confirmed by the parameter distribution in these simulation types of relatively high k_4_ (Fig. S3B, Table S5). We further confirm that higher membrane dimer levels are associated with increased fendiline concentration for all of the simulations (Fig. 6, Table S19-S20). Taken together, these observations confirm our hypothesis that fendiline treatment can counterintuitively result in increased VLP production when filament growth is slow (characterized by low k_3_ and k_3,1_’) or VLP budding rate constant is high (characterized by high k_4_).

**Figure 5.**
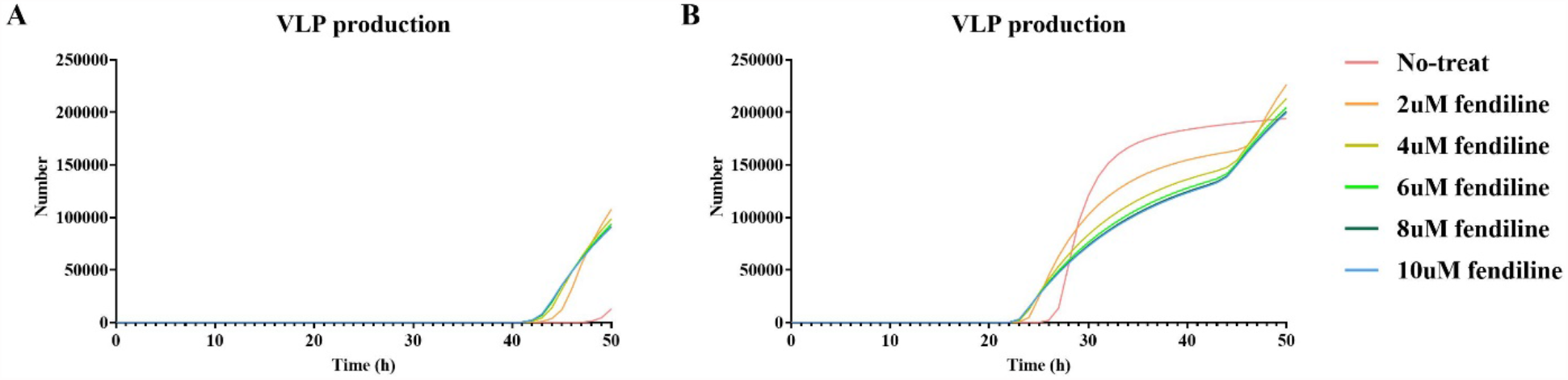
Dynamic of VLP production in two types of fendiline-induced VLP increase groups. Higher VLP production at 48h with fendiline treatment due to (A) late VLP production time or (B) fluctuation.

**Figure 6.**
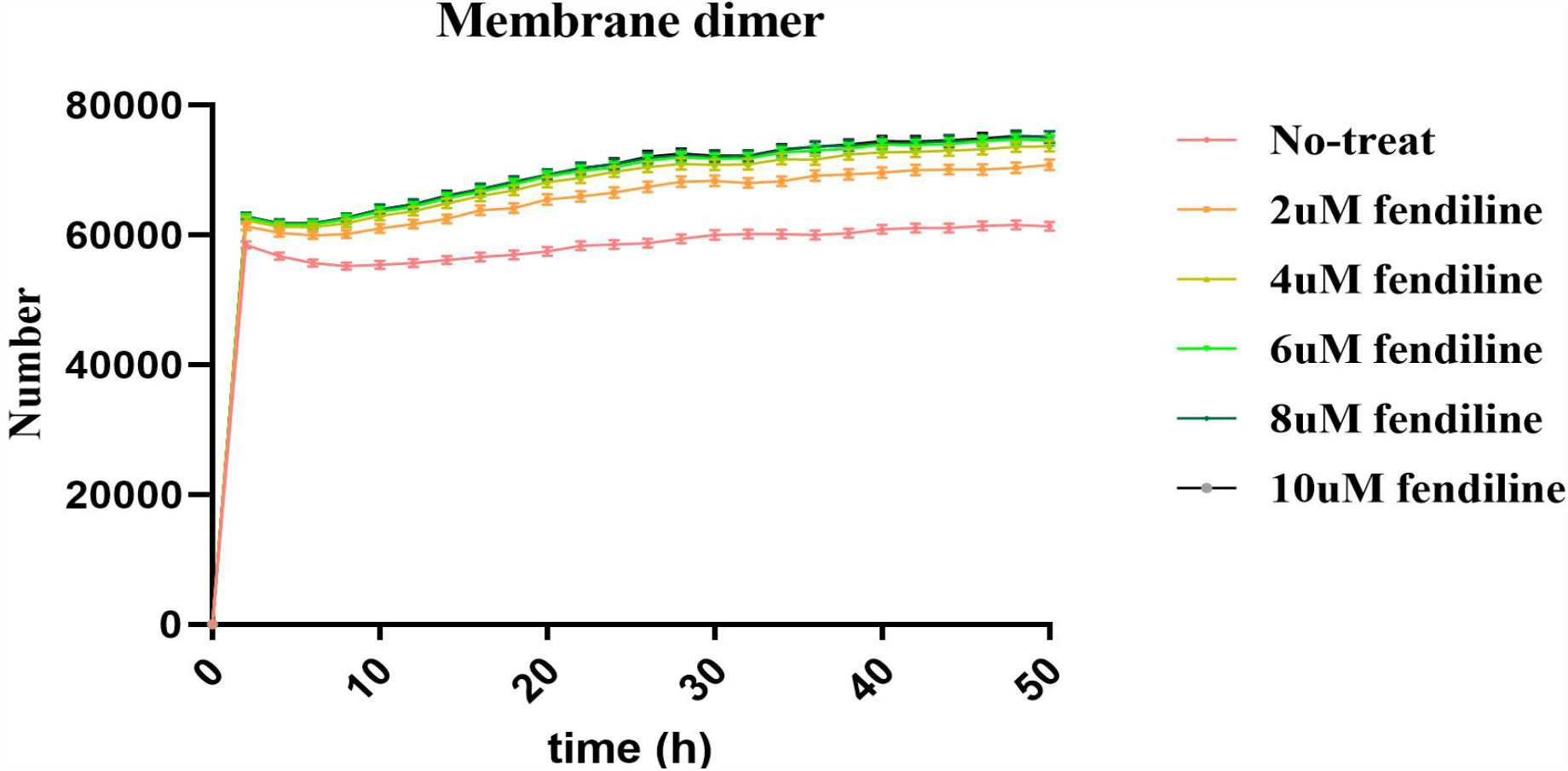
Concentration of membrane dimer in fendiline-induced VLP increase simulation. Higher concentration of fendiline will bring higher concentration of membrane dimer pool. Error bars indicate SEM. The figure is plotted based on every two hours.

Also, in all 199 parameter sets having elevated VLP production with 2 μM fendiline treatment under 24 h, only 58 of them still have increased VLP production at 48 h (Table S4). This suggests that the most fendiline induced increase in VLP production will be dissipated over the course of treatment. It means that though slow filament growth may be the major reason for the fendiline induced increase in VLP production, the phenomena will only be detectable around the budding start time and will not persist into longer-term effect. On the other hand, while high VLP budding rate constant may result in fendiline-induced VLP production during any time in VLP budding stage, they are rare cases due to random fluctuations at the specific measurement time, and cannot be broadly considered as “fendiline has increased VLP production”.

Based on our analysis and experimental data, we conclude that fendiline can decrease VLP production rates but could also result in earlier budding start times, which may lead to increased VLP production around the VLP budding start time. However, in a longer-term, fendiline should still decrease VLP production effectively. Though the effect may be weakened by high VLP budding rate, it will not be reversed. Thus, we next evaluate fendiline as a potential treatment for EBOV infection in most situations, and the conclusion should not be affected by the observed fendiline induced VLP production at some measurement time points.

### Delayed usage of fendiline shows that it can be an effective drug even after budding has already started

Fendiline affects VLP production in two ways: (1) It reduces the VLP production rate but (2) results in earlier VLP production time. The first impact depends on fendiline’s effect on the VLP budding step. The second impact depends on its influence on VP40 filament growth, which is important prior to the VLP budding stage. Thus, our next question is: what is the impact of fendiline application time on VLP production? To test the impact of infection stage on fendiline efficacy, we simulated 2 μM and 10 μM fendiline treatment starting at 0, 12, 24 and 36 hours post infection, and recorded VLP production at 24 h and 48 h post infection. Since the concentration of fendiline is constant in our study, the earlier fendiline is used, the more VLP reduction is achieved as expected (Fig. 7A-B). However, when we looked at the average VLP reduction per hour, it is obvious that the latter fendiline is applied, the higher the efficiency (Fig. 7C-D, Table S6, S9). This suggests that fendiline is relatively more effective later in the infection cycle.

**Figure 7.**
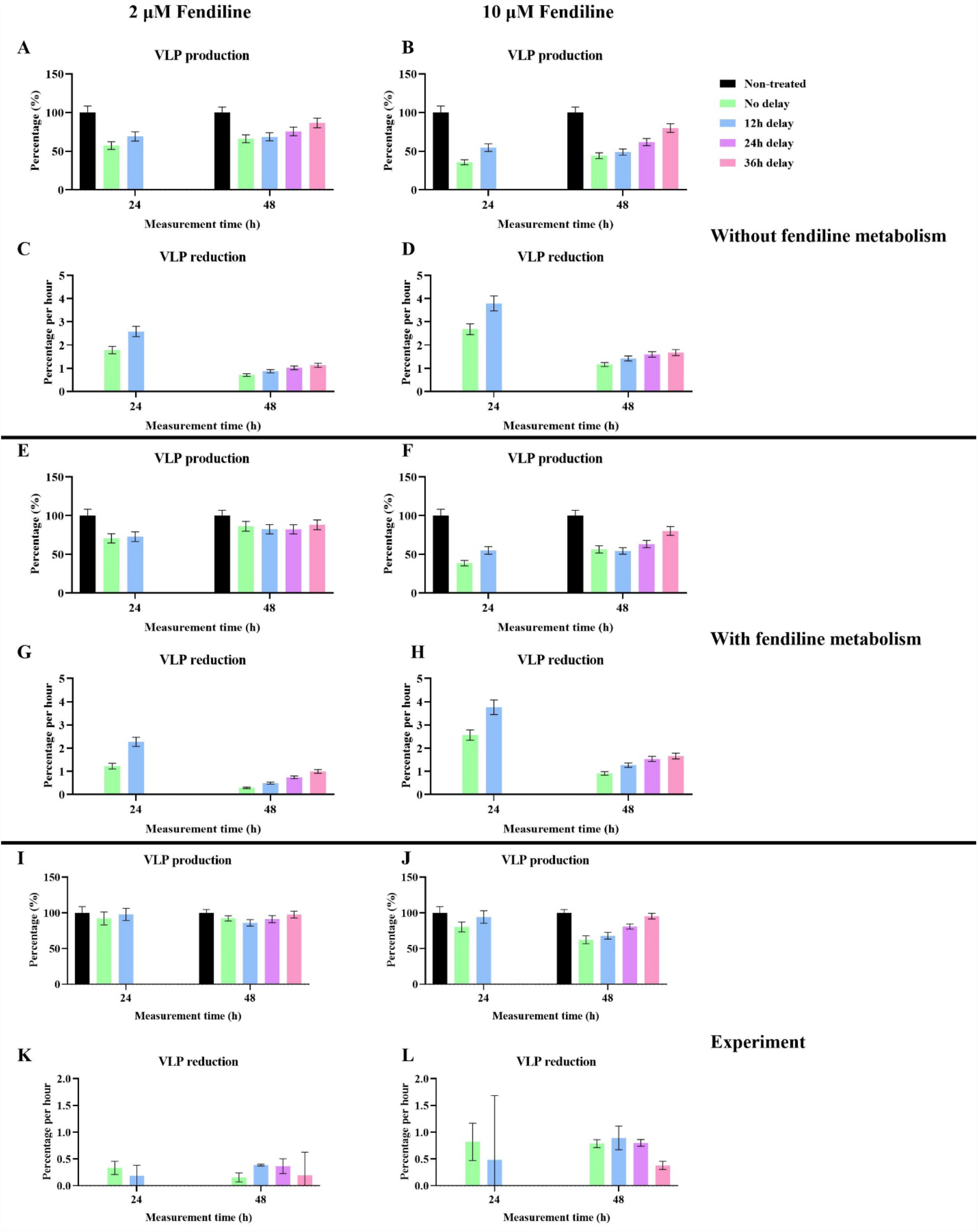
VLP inhibition by different application time of fendiline. (A) Simulation of VLP production under 2μM fendiline applied at 0, 12, 24 and 36 hours post infection without the metabolism of fendiline. (B) Simulation of VLP production under 10μM fendiline applied at 0, 12, 24 and 36 hours post infection without the metabolism of fendiline. (C) Simulation of VLP reduction percentage per hour under 2μM fendiline applied at 0, 12, 24 and 36 hours post infection without the metabolism of fendiline. (D) Simulation of VLP reduction percentage per hour under 10μM fendiline applied at 0, 12, 24 and 36 hours post infection without the metabolism of fendiline. (E) Simulation of VLP production under 2μM fendiline applied at 0, 12, 24 and 36 hours post infection with 20 h half-life of fendiline. (F) Simulation of VLP production under 10μM fendiline applied at 0, 12, 24 and 36 hours post infection with 20 h half-life of fendiline. (G) Simulation of VLP reduction percentage per hour under 2μM fendiline applied at 0, 12, 24 and 36 hours post infection with 20 h half-life of fendiline. (H) Simulation of VLP reduction percentage per hour under 10μM fendiline applied at 0, 12, 24 and 36 hours post infection with 20 h half-life of fendiline. (I) Experiment of VLP production under 2μM fendiline applied at 0, 12, 24 and 36 hours post infection. (J) Experiment of VLP production under 10μM fendiline applied at 0, 12, 24 and 36 hours post infection. (L) Experiment of VLP reduction percentage per hour under 2μM fendiline applied at 0, 12, 24 and 36 hours post infection. (L) Experiment of VLP reduction percentage per hour under 10μM fendiline applied at 0, 12, 24 and 36 hours post infection. Error bars indicate the SEM.

Since the half-life of fendiline in plasma is about 20 h (32), and we have concluded that fendiline is relatively more effective in later applications under constant concentration, we wanted to simulate a case of single dosage of fendiline and see if a best application time exists when considering the effect of pharmacokinetics. Another round of simulation was conducted with fendiline decay rate of 9.625×10^−6^/s (calculated based on 20 h half-life). Because the half-life of fendiline is 20 h, we extended the simulation time to 74 h. The result shows that the fendiline inhibition on VLP production is stronger when the application time is later. However, when fendiline is applied too late there might be reduced effects due to reduced exposure time before the end of the simulation (Fig. 7E-F, Table S7, S9). Again, when we looked at the average VLP reduction per hour for different fendiline application times, the efficiency will increase with later usage of fendiline (Fig. 7G-H, Table S7). To make sure this is not due to fluctuations, we also looked at the average VLP production dynamics of the different fendiline application times (Fig. 8). We can see that the production will be inhibited for a period of time after the drug is applied, which is in line with PS concentration reduction (Fig. S4). And the later the drug is applied, the more overall reduction it will have in the longer term.

**Figure 8.**
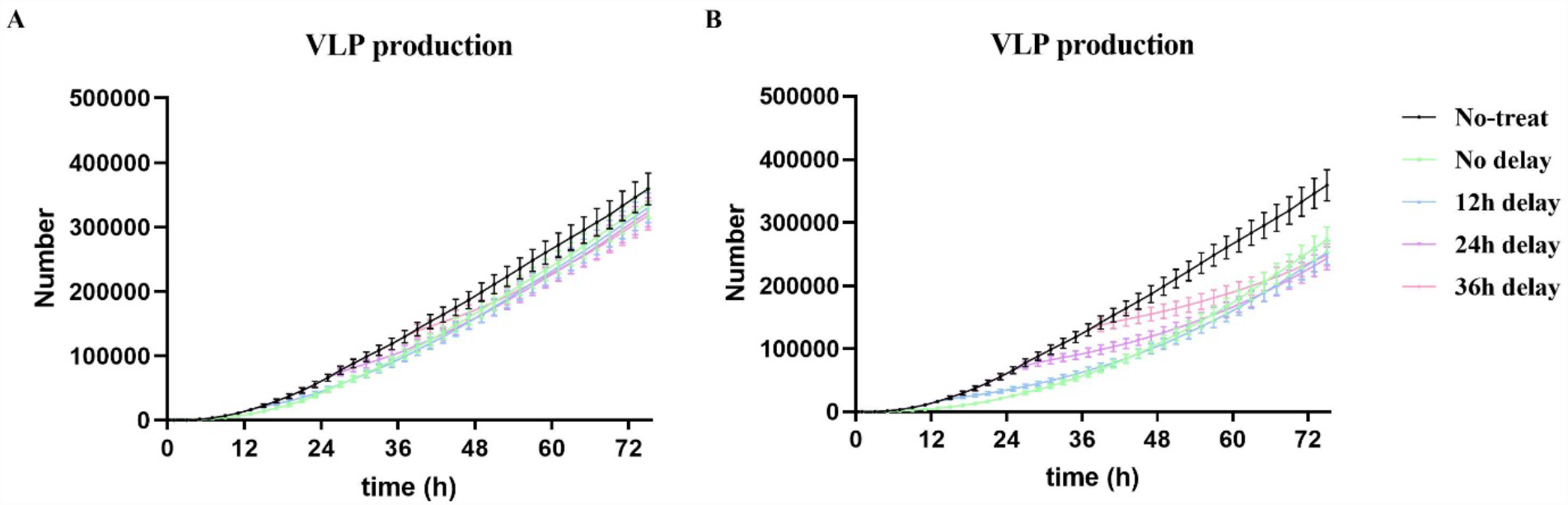
VLP production dynamic of one-dose fendiline simulation. (A) 2μM of fendiline applied at 0, 12, 24 and 36 hours post infection. (B) 10μM of fendiline applied at 0, 12, 24 and 36 hours post infection. Error bars indicate the SEM. The figure is plotted based on every two hours.

To confirm our findings in cells expressing VP40, we expressed EGFP-VP40 as previously described (15, 17) and treated cells with fendiline or vehicle control (DMSO) at different time points (0, 12, 24, and 36 hours post-transfection). The pre-VLP (VLPs localized around the cell membrane) formation was then assessed at 24 h and 48 h using fluorescence microscopy. As pre-VLPs are the precursor of VLPs, we believe that the experimental trend in pre-VLP changes can represent the simulation trend in VLP change. The experimental results show that VLP production is increased with later usage of fendiline for most cases (Fig. 7I-J, Table S8), which is the same as a constant concentration of fendiline application (Fig. 7A-B). However, when 2μM fendiline application changes from 12 h delay to 24 h delay, the VLP production is decreased at 48h (Fig. I, Table S8), which reproduces the simulation results when considering the pharmacokinetics of fendiline (Fig. 7E). This likely indicates that in cell culture there might be a far slower clearance of fendiline compared to human body, and the concentration of fendiline remains almost constant.

When calculating the VLP reduction per hour from the experiments, increased VLP reduction per hour is observed at 48 h when fendiline application is delayed from 0 h to 12 h for both 2μM and 10 μM fendiline (Fig. 7K-L), which aligns with our simulations (Fig. C-D, G-H). From this result, we believe the experimental data confirms our predictions that fendiline is relatively more efficient when applied at later times within the viral life cycle. However, we also observe that the VLP reduction per hour is decreased in other fendiline application times. This is possibly caused by the fact that fendiline-induced PS reduction may be slower in experimental conditions compared to our simulations, as our model has not considered the fendiline absorption process and how fast PS cycling is happening. As a result, when the application time of fendiline is close to measurement time (12-24 h prior), the real “effective treatment time” of fendiline could be much shorter than assumed in the simulations. This can be proved by our finding that when fendiline is applied 12-24 h prior to the measurement time, the difference between 2μM and 10 μM fendiline is not significant (Fig. S5, Table S10), indicating fendiline may not have enough time to be effective.

From these results, we conclude that when viral budding is already established and mature within a single cell, application of fendiline will be relatively more effective. Thus, fendiline can be a useful treatment for cells in the egress stage of EBOV infection on a single cell level. But we need to be careful, as there can be an innate delay from application time and effective time that should potentially be explicitly included in future model iterations. Also, it remains unclear how these single cell dynamics would affect the overall efficacy of fendiline in a population of cells that could all be in different stages of infection, which is outside the scope of the current work, but multi-scale modeling efforts are under way to answer this question.

### Co-treatments simulation identified extra beneficial effect of fendiline with certain step-targeted treatments and high viral protein expression mutant strain

Our final question is how fendiline could work with other treatments targeted to specific steps in the VP40 viral matrix assembly process. We have hypothesized that some of the step-targeted treatments may have synergistic effects with PS-targeted treatment by PRCC in a prior study (14). We further explored this here by changing target parameters (r_1_, k_1_, k_2_, k_3_, k_4_) to half while maintaining the values of other parameter values in the fendiline treatment simulations. Here we focus on 2 μM fendiline since it is closer to the expected therapeutic plasma concentration (32). To determine if potential combination treatments are synergistic, we compare the simulated co-treatment efficiency with the product of their individual efficacies (representing additive effects).

First, simulation of k_4_ (VLP budding rate constant) targeted treatment shows a significant synergistic effect with fendiline. On average, 3.65% and 7.46% additional treatment effects are achieved at 24h and 48h, separately, compared to an additive treatment effect (Fig. 9A, Table S11). This is expected since k_4_ is more rate-limiting and thus increases the regulation effect of PS when the step is targeted. The conclusion is also statistically supported by t-tests (Table 2).

**Table 2.** T-test for co-treatments simulation.

|  | 24h |  | 48h |  |
| --- | --- | --- | --- | --- |
|  | Average additional reduction | p-value | Average additional reduction | p-value |
| Fendiline with $r_1$-targeted | 1.83% | 0.0001 | 1.37% | <0.0001 |
| Fendiline with $k_1$-targeted | -0.04% | 0.5742 | -0.02% | 0.6049 |
| Fendiline with $k_2$-targeted | -0.73% | 0.1208 | -0.15% | 0.2141 |
| Fendiline with $k_3$-targeted | -0.07% | 0.7546 | -1.21% | <0.0001 |
| Fendiline with $k_4$-targeted | -3.65% | <0.0001 | -7.46% | <0.0001 |

**Figure 9.**
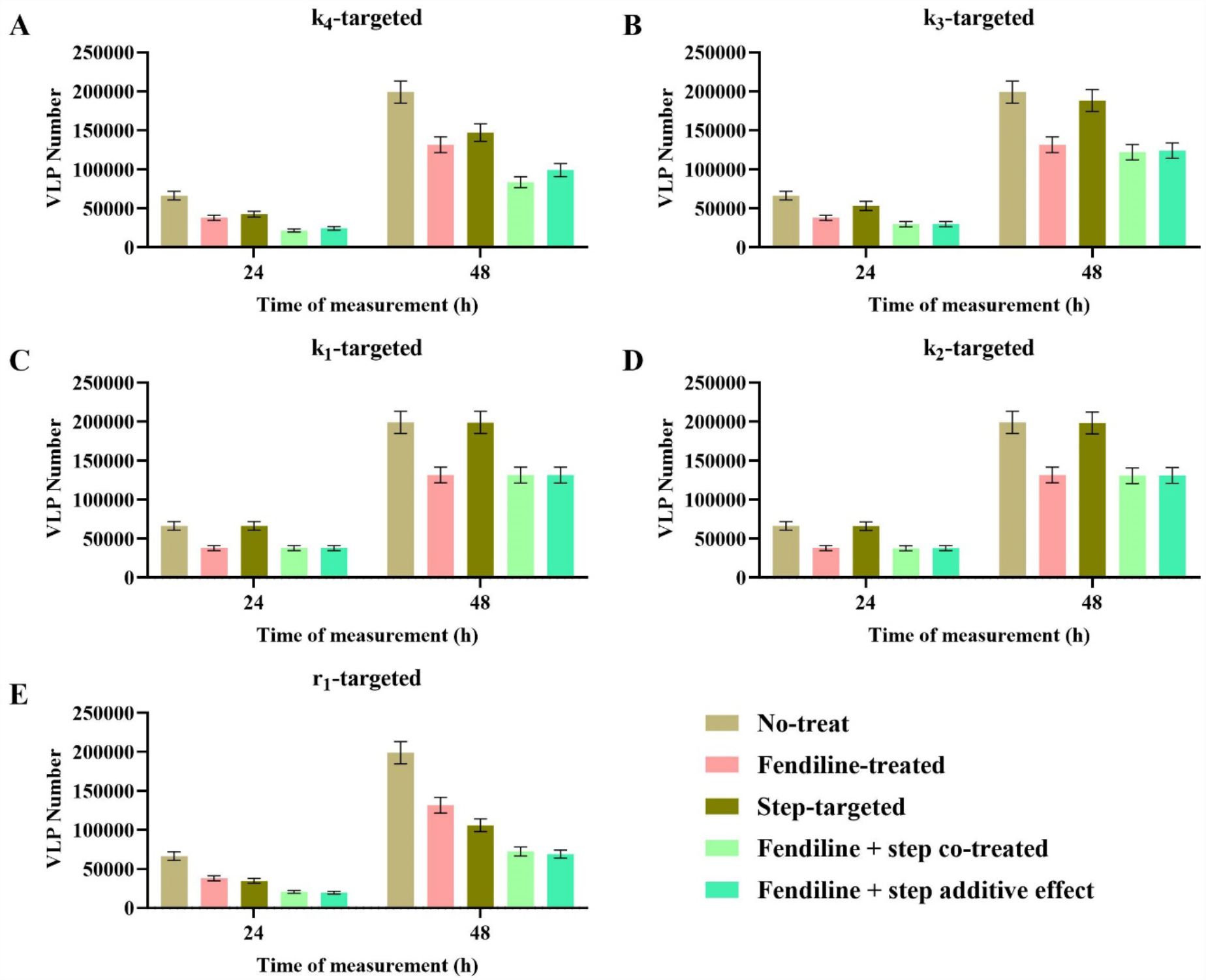
VLP production of step-targeted and fendiline co-treatment. Co-treatment of fendiline with (A) k_4_ shows an obvious synergistic effect; with (B) k_3_ shows a weak synergistic effect at 48h; with (C) k_1_ and (D) k_2_ shows independent effect; with (E) r_1_ shows an antagonistic effect. Error bars indicate the SEM.

Simulation of k_3_ (filament growth forward rate constant) targeted treatment shows statistically significant synergy with fendiline at 48 h (1.21%), but not at 24 h (Fig. 9B, Table 2, S12). The reason can be inferred from our earlier finding regarding the impact of k_3_ on timing of budding start (Fig. 5A, S3A). If k_3_ is targeted and lowered, the budding time of VLP will be postponed. The co-treatment of fendiline will counter-act this change and make the production of VLP higher around the budding start time. This mechanism of fendiline counter-acting the impact of a lower k_3_ value would be more obvious in short-term (24 h), as it is closer to the budding start time. But for longer-term, they should still have a steady synergistic treatment efficiency.

Simulation of k_1_ (dimerization forward rate constant) and k_2_ (membrane association rate constant) targeted treatment shows that they hardly impact VLP production on their own, and they are independent of fendiline treatment (Fig. 9C-D, Table 2, S13, S14). The reason that they are not synergistic could be that these steps are not rate-limiting in our system.

Finally, simulation of r_1_ (VP40 production rate) targeted treatment shows an antagonistic efficiency with fendiline (Fig. 9E, Table S15). P-values from t-test show that the efficiency of co-treatment is always lower than the product of individual treatments (Table 2). But this does not mean we cannot apply those two kinds of treatments simultaneously. They can be used as the efficiency of co-treatment is always higher than any single usage. This only suggests that there are no synergistic effects. On the other hand, it means that fendiline can be more useful when r_1_ is higher, which could represent a viral strain with high viral protein expression.

## Conclusion

Fendiline has been proposed to be a potential treatment for EBOV infection (9). Here, we explore mechanisms, dynamics, and potential co-treatments of fendiline with a computational model of VP40 VLP assembly and budding.

Our model is developed from a previously published version (14), and incorporates the latest knowledge of VP40 budding and the nucleation-elongation theory. Our findings suggest that the filament growth of VP40 follows the nucleation-elongation process. While this process has not been specifically studied in VP40 oligomerization, it is widely accepted as a mechanism in biopolymer assembly (20, 33). We also find that higher PS may decrease the difference in filament stability between the nucleation and elongation processes. Since nucleation is usually slow and rate-limiting (20), we believe it to be a good target for interrupting the VP40 assembly pathway.

We hope the nucleation-elongation process can be tested experimentally for VP40 in the future, as it can improve our understanding of the assembly process of VP40 and help evaluate new therapies that target VP40 oligomerization.

Our simulations indicate that fendiline can effectively suppress the production of VLPs in most cases, while fendiline can also increase the production of VLPs for specific parameter sets at certain time points. This dual effect is related to the fact that as fendiline decreases the concentration of cell membrane PS, the cell membrane dimer pool of VP40 will be enlarged and serve as a reservoir which will promote the maturation of growing filaments and bring the budding time of VLPs earlier. Increasing VLP production in response to fendiline treatment happens around VLP budding start time, and should be disappearing with longer time treatment. Due to the existence of fluctuation in VLP production, the shift in VLP production start time by fendiline treatment may also cause higher VLP production under fendiline treatment at some time points, especially when the VLP budding rate is high. However, from our analysis, neither of these “fendiline induced VLP production” cases are persistent in the longer-term. Experimental data that support this computational finding (15), indicates that fendiline can effectively reduce VLP production in the longer-term in the VP40 system. Also, as our model simulates at a single cell level, the increased VLP production may be averaged out by the whole asynchronous population, and the effect of fendiline at multi-cell level will be further evaluated in future computational studies.

Since fendiline may result in earlier VLP production, we also find that the treatment efficiency of fendiline is higher when the application time is later. When we consider the dynamics of fendiline metabolism and the application of fendiline with some delay after infection, our simulation indicates that fendiline might be more effective in cells where VLP budding stage has already been established, since it can suppress the VLP budding directly. On the other hand, it doesn’t mean that fendiline cannot be applied at early stages. Most of our simulations still show reduced VLP production when fendiline is applied at early time points (Fig. 3), and the increased VLP production may be averaged out over a large population of cells that are all in different stages of infection. While our current model cannot confirm this notion, we are developing multi-scale models to study the effect of this cellular stochasticity on tissue-scale outcomes in the future.

We also explore the co-treatment of fendiline with other hypothesized step-targeted treatment in the VP40 assembly and budding pathway. We want to pay extra attention to k_3_-targeted treatment, since graphene is proposed to be an inhibitor to the filament growth process (34). The co-treatments will have a synergistic effect in the longer term. But for the short-term, since k_3_-targeted treatment can slow down the filament maturation, it may lead to increased VLP production under fendiline treatment in some cases. These cases are characterized by a strong treatment efficiency of k_3_-targeted treatment alone (Table S12). Due to this and the previous finding that fendiline is relatively more effective at later times, we believe that if we are going to use both fendiline and k_3_-targeted therapy as treatment, it may be better to apply k_3_-targeted treatment first and determine the application time of fendiline based on the treatment efficiency of the k_3_-targeted drug.

Fendiline also shows a strong synergistic effect with k_4_-targeted treatments, suggesting that it will be extra beneficial to apply the co-treatments and achieve better efficiency. Though for r_1_-targeted treatments, co-treating with fendiline does not show synergistic effects, they can also be used since the adding of fendiline is still better than single-treatment. Moreover, the simulation also informs us that fendiline can be extra useful when r_1_ is high, which indicates a high expression (more viral proteins) EBOV mutant strain (35).

While we are only evaluating fendiline in our study, our model can be used to evaluate other potential PS-targeted or VP40-targeted treatments, such as staurosporines (36, 37). A recent study found that sangivamycin, a protein kinase C inhibitor, can interrupt VP40 membrane association and decrease VLP production, and proposes it to be a new EBOV therapy (38). Our model may also support the evaluation of sangivamycin, as its influence on the VP40 profile is similar to fendiline.

There are also limitations to our study. Our simulation is based on cell culture, and only represents infection on a single cell-level. No intercellular infection exists in our model. There are also differences between the VP40 system and authentic EBOV for which our current model does not account. Due to these limitations, we do not aim to make clinical suggestions, but consider this work a step toward improved mechanistic understanding and drug development.

Overall, we have evaluated the impact of fendiline on VP40 VLP production with our model simulation. Though experimental studies generally propose that fendiline is effective in suppressing VLP production, we further explore the potential VLP production increase in short time, the efficiency in different time stages and co-treatment effects with other hypothesized VP40 assembly-targeted treatments. The dual effect of fendiline makes the case more complicated, but our results indicate that in general, fendiline has potential to reduce VP40 VLP production. It can still be effective if the treatment is delayed, and work well with other step-targeted treatments, and be particularly effective against EBOV strains with high viral protein expression.

## Methods

### ODE-based model construction

Our model is updated from the “Budding” model constructed in our last work (14). Model updates that are new in this work are outlined here. According to the latest knowledge on VP40 filament growth (19), we alter the building block from cell membrane hexamer to dimer. Also, we replace the “filament stabilization” mechanisms from the prior work with a direct nucleation-elongation process for VP40 oligomerization (Fig. 10A). ‘As1’-’As4’ models are constructed to determine the PS influence on the process (Table 1). We also include a simple PS metabolism network to divide our PS pool into cytoplasmic and cell membrane compartments (Fig. 10B).

**Figure 10.**
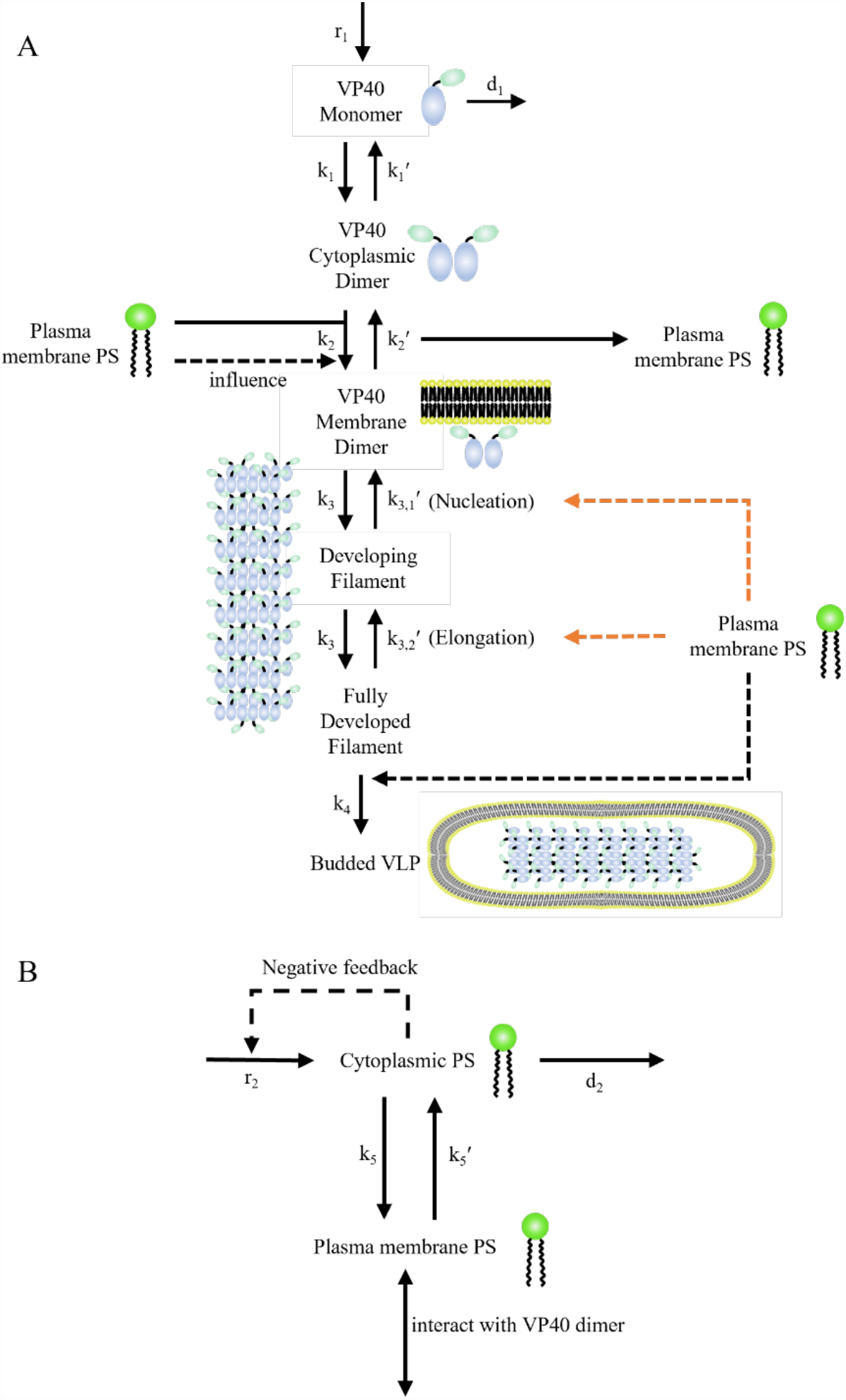
Diagram of the VP40 model. (A) VP40 production, assembly, and budding process. (B) PS metabolism network. All black lines are known reactions while all orange lines are hypothesized mechanisms proposed in this study. Solid lines are direct interactions while dashed lines are influence. Influence of PS on nucleation-elongation process is tested in ‘As1’-’As4’ models (Table 1).

ODEs for the main processes are presented in Eq. (1)-(10).

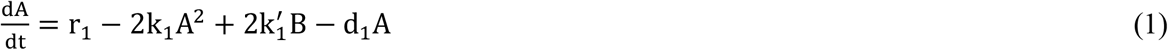

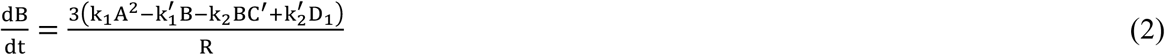

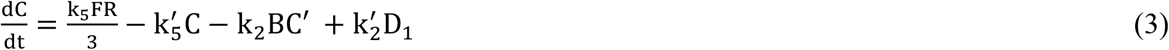

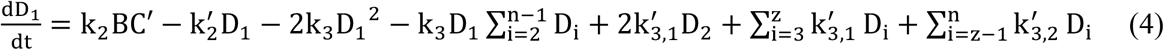

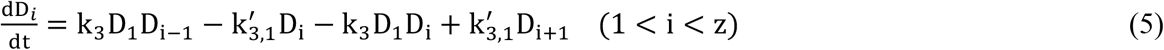

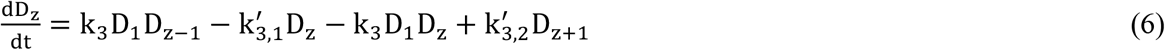

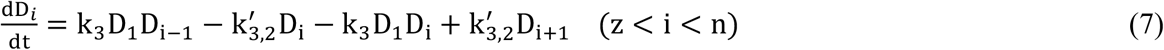

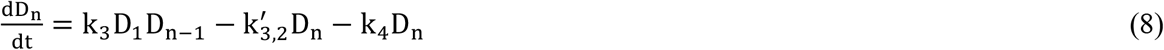

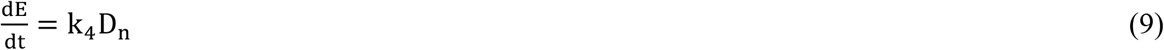

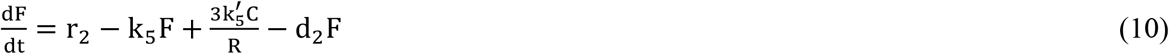

Initial conditions:

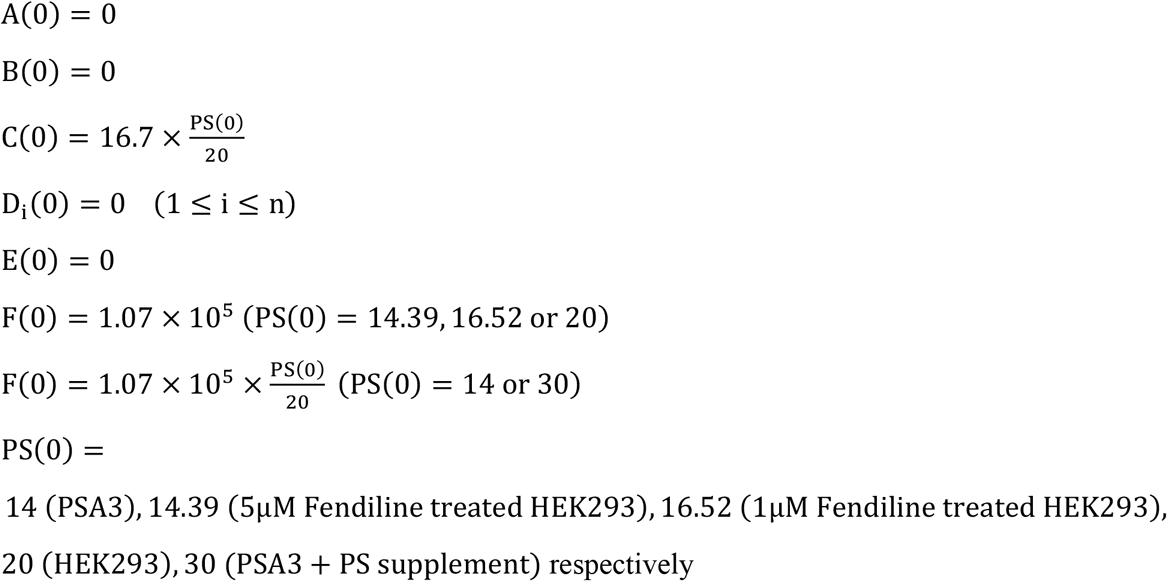

A: VP40 monomer in cytoplasm (nM).

B: VP40 dimer in cytoplasm (nM).

C: Plasma membrane phosphatidylserine (nmol/dm^2^).

C′: Plasma membrane Phosphatidylserine available to interact with cytoplasmic VP40 dimer (nmol/dm^2^).

D_i_: Developing matrix protein consists of i VP40 dimers (nmol/dm^2^).

i: Number of dimers in developing filament.

z: size of oligomer where the reverse rate constant change from k_3,1_′to k_3,2_′.

n: Number of dimers in a mature filament. n = 2310 in our model.

E: Budded VLP (nmol/dm^2^).

F: Cytoplasmic phosphatidylserine (nM)

PS: Plasma membrane phosphatidylserine (%).

PS level will be updated by the concentration of C through Eq. (11).

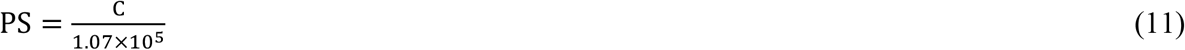

The adding of R (radius of cell, dm) is used to compensate the unit change between surface (nmol/dm^2^) and volume (nM) concentration.

The values and calculations of all parameters are listed in Table S3.

### Influence of PS on VP40 budding system

In our model, VP40 dimer membrane association and VLP budding steps are directly influenced by cell membrane PS level as described in our previous study. Equations are updated to fit the current model and avoid negative values in Eq. (12)-(14).

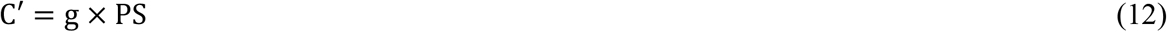

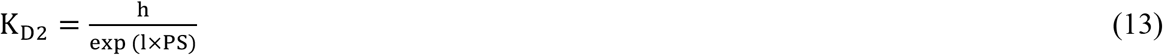

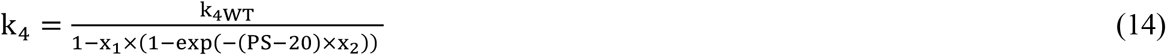

Fitted values of g, h, l, x_1_ and x_2_ are included in Table S3.

The influence of PS on nucleation-elongation process is tested in ‘As1’-’As4’ models (Table 1), which is represented in Eq. (15)-(16).

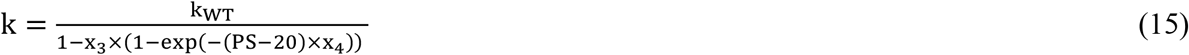

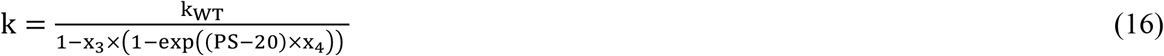

k: involved parameter k_3,1_′ (‘As1’, ‘As2’ model) or k_3,2_′ (‘As3’, ‘As4’ model) changing with PS level.

k_WT_: involved parameter k_3,1_′ (‘As1’, ‘As2’ model) or k_3,2_′ (‘As3’, ‘As4’ model) under 20% PS level.

Values of parameter x_3_ and x_4_ can be found in Table S3.

### Influence of PS on its production

Negative feedback exists in PS production, where a high PS level will lead to a lower PS production rate (39–41). To reflect this negative feedback, we assume that the inhibition is caused by cytoplasmic PS concentration, apply a hill-like function in the production of PS as Eq. (17).

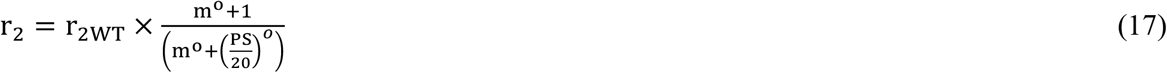

The data used for fitting is shown in Table S16. Fitted values of parameter m and o can be found in Table S3.

### Influence of PSA-3, fendiline and PS supplement on PS network in calibration

As PS metabolism network is included in our model, experimental scenarios will be linked to a specific step in the network. In previous studies, it has been shown that fendiline will inhibit the activity of acid sphingomyelinase (ASM), decrease the hydrolysis of sphingomyelin, elevate sphingomyelin levels, and block the recycling of PS to cell membrane (10, 11). We reflect this mechanism as an influence on the membrane association constant of PS.

The ratio of plasma membrane PS to cytoplasmic PS is 6 under WT situation (36). Under steady state situation, when no VP40 exists in the system, we will have Eq. (18)-(19):

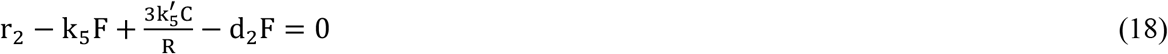

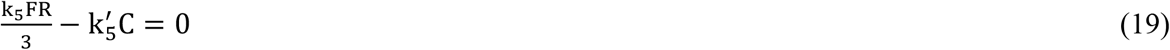

Combining the two equations will get Eq. (20):

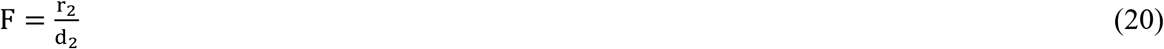

While fendiline does not affect production or decay of PS, cytoplasmic PS level remains the same. From Eq. (19), Eq. (21) can be deducted:

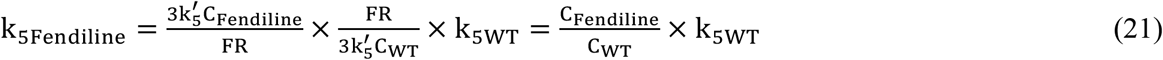

The values of k_5_ under 1 µm and 5 µm fendiline is calculated accordingly (Table S3, S17).

PSA-3 cells are genetically compromised in PS production (13, 42), and we apply this influence on r_2_. While PSA-3 does not affect the localization of PS, from Eq. (19), Eq. (22) can be conducted:

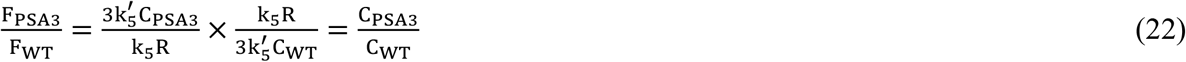

Combining Eq. (20) and Eq. (22) will get Eq. (23):

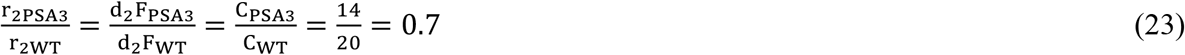

Since PS production is regulated by cytoplasmic PS concentration, Eq. (17) needed to be considered, and the final value of PSA-3 PS production rate will be calculated by Eq. (24):

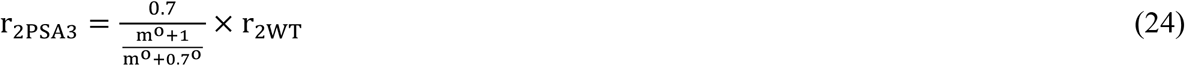

Supplement of PS will be reflected in PS production as well. Applying Eq. (23) will get Eq. (25):

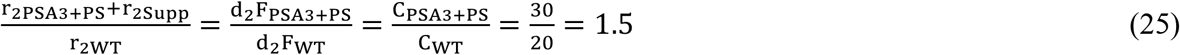

Supplement of PS is regarded as a constant number not affected by cytoplasm PS level, and the innate PS production ability in PSA3 with PS supplement group should be the same as PSA3 group, thus the supplement of PS is calculated accordingly in Eq. (26).

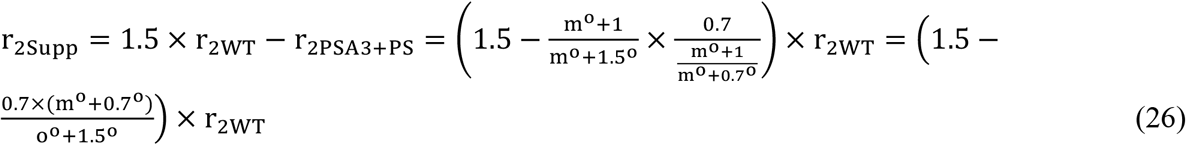

### Parameter estimation and calibration

Latin hypercube sampling (LHS) is used to sample the parameters within the ranges given in Table S3. The sampling for x_2_ and z is on a linear-scale, while for other parameters it is on a log scale. The top 5 parameter sets with lowest cost are used to analyze the influence of PS on the nucleation-elongation process. In fendiline simulations, all parameter sets with a cost ≤ 3 or score ≥ 5 are used for analysis to reflect individual differences and account for parameter uncertainty. A diagram of the parameter estimation process is shown in Fig. 11. The cost represents the average fold change in prediction to data under each type of data as listed in Eq. (27):

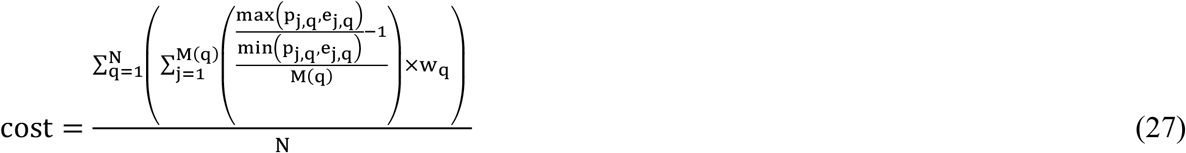

N: Number of different data type

M(q): Number of data in the qth data type

e_j,q_:jth experiment data in the qth data type

p_j,q_:jth model prediction in the qth data type

W_q_:weight assigned to qth data type (Table S18)

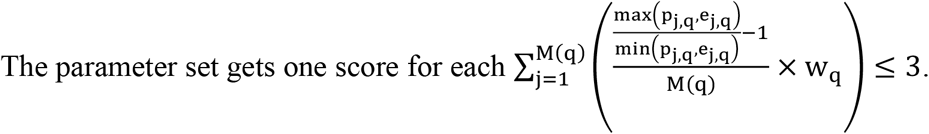

Any predictions at a certain time point are calculated from the average values of ± 2 h around prediction time (e.g., 22-26 h for 24 h) to avoid potential extreme values from fluctuation. The application of average prediction, cost and score will decrease the bias from fluctuation and a single data type and explore the system with higher diversity. Prediction of relative oligomer frequency on each size of oligomer is calculated from the average values within 1 oligomer size around prediction size (e.g., tetramer-octamer for hexamer) except for dimer.

**Figure 11.**
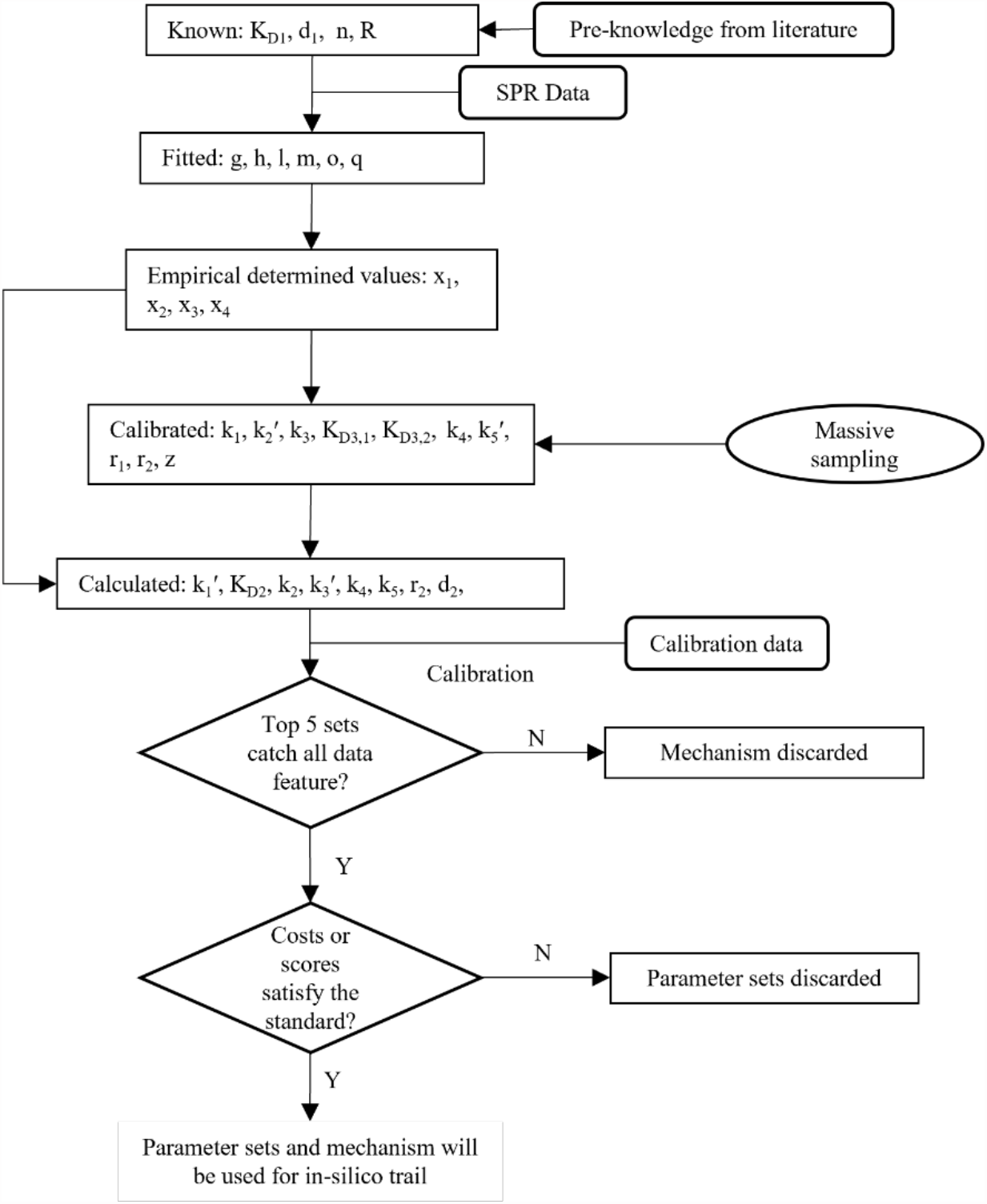
Parameterization process for the model. Top 5 lowest cost samples from each model will be analyzed, and the model is be discarded if all the data features are not caught. Rest of the models with parameter sets that pass the requirement will be used for further in-silico trail.

Data used for calibration is the same as those in our previous work (14), with updated interpretations:

- VP40 oligomer ratio is defined as the ratio of VP40s amount in 6-48mer to those in monomer and dimer.
- Relative oligomer frequency is defined as the relative oligomer amount from hexamer to 42mer to the sum of them.

Other definitions remain the same:

- VLP production is defined as the number of VLP produced by a single cell.
- VP40 budding ratio is refined as the ratio of VP40s amount in budded VLPs to those in cell.
- VP40 plasma membrane localization is defined as the ratio of VP40s amount in membrane dimer-48mer to those in cytoplasm monomer and dimer.

### Application of in silico fendiline simulation

Fendiline is applied as input to the system in simulation with the feasible parameter sets and models identified in the calibration.

The relationship between fendiline and k_5_ is fitted to empirical Eq. (28) with the k_5_ value under 1, 5 or 10 µm of fendiline (Table S17).

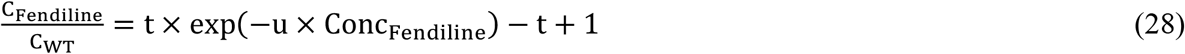

Fitted values of u and t are listed in Table S3. Combining Eq. (21) and Eq. (28), k_5Fendiline_ will be calculated according to Eq. (29) in the simulation:

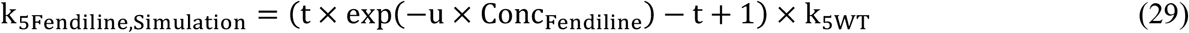

In co-treatment with other step-targeted treatments, the rate-constant for the targeted treatment step is reduced to half while others remain the same for simulation.

During simulations, VLP production lower than 1 will be considered 0.

### Experimental cellular studies with fendiline

HEK293 cells were maintained and transfected with EGFP-VP40 plasmid DNA as previously described (15) in DMEM containing 10% FBS and 1% penicillin/streptomycin. Transfections were done in DMEM containing 10% FBS in the absence of penicillin/streptomycin. Cells were maintained in DMEM containing 10% FBS following transfections and were treated with either vehicle (DMSO) or fendiline (at varying concentrations in DMSO) for analysis at different time points (24 or 48 hours). Confocal imaging was performed on a Nikon Eclipse Ti Confocal microscope (Nikon Instruments, Melville, NY) using a 60x 1.4 numerical aperture oil objective (or a 100x 1.45 numerical oil objective as needed) or a Zeiss LSM 710 using a 63x 1.4 numerical aperture objective. Image analysis (plasma membrane localization pre-VLP formation) was performed by counting pre-VLPs at the plasma membrane per cell slice by scanning the Z plane of the image. The number of preVLPs were assessed per imaging frame for an equal number of VP40 expressing cells over the course of three independent experiments.

## Data availability

All model code is available on Zenodo under DOI 10.5281/zenodo.7921784. Add data used to generate figures are available in the supplemental material.

## Funding acknowledgements

This project was funded with support from the Indiana Clinical and Translational Sciences Institute which is funded in part by Award Number UM1TR004402 from the National Institutes of Health, National Center for Advancing Translational Sciences, Clinical and Translational Sciences Award (to EP and RVS); and AI081077 (to RVS); and T32GM075762 (to MH). The content is solely the responsibility of the authors and does not necessarily represent the official views of the National Institutes of Health. This material is based upon work supported by the National Science Foundation under Grant No. 2143866 (to EP). This work used the Extreme Science and Engineering Discovery Environment (XSEDE), which is supported by National Science Foundation grant number ACI-1548562. Anvil at Purdue was used through allocation TG-MDE220002 (to EP).

## References

1. Feldmann, H., and Geisbert, T. W. (2011) Ebola haemorrhagic fever. The Lancet. 377, 849–862

2. Feldmann, H., Jones, S., Klenk, H. D., and Schnittler, H. J. (2003) Ebola virus: From discovery to vaccine. Nat Rev Immunol. 3, 677–685

3. Ebola Virus Disease Distribution Map: Cases of Ebola Virus Disease in Africa Since 1976 | History | Ebola (Ebola Virus Disease) | CDC (2021) [online] https://www.cdc.gov/vhf/ebola/history/distribution-map.html x(Accessed August 10, 2022)

4. Aschenbrenner, D. S. (2021) Monoclonal Antibody Approved to Treat Ebola. American Journal of Nursing. 121, 22

5. Mulangu, S., Dodd, L. E., Davey, R. T., Tshiani Mbaya, O., Proschan, M., Mukadi, D., Lusakibanza Manzo, M., Nzolo, D., Tshomba Oloma, A., Ibanda, A., Ali, R., Coulibaly, S., Levine, A. C., Grais, R., Diaz, J., Lane, H. C., Muyembe-Tamfum, J.-J., and the PALM Writing Group (2019) A Randomized, Controlled Trial of Ebola Virus Disease Therapeutics. New England Journal of Medicine. 381, 2293–2303

6. Markham, A. (2021) REGN-EB3: First Approval. Drugs. 81, 175–178

7. Bayer, R., and Mannhold, R. (1987) Fendiline: a review of its basic pharmacological and clinical properties. Pharmatherapeutica. 5, 103—136

8. Fendiline | C23H25N - PubChem [online] https://pubchem.ncbi.nlm.nih.gov/compound/Fendiline#section=Chemical-Vendors x(Accessed August 10, 2022)

9. Husby, M. L., and Stahelin, R. (2018) Repurposing Fendiline as a novel anti-viral therapeutic. The FASEB Journal. 32, 671.9–671.9

10. Cho, K., van der Hoeven, D., Zhou, Y., Maekawa, M., Ma, X., Chen, W., Fairn, G. D., and Hancock, J. F. (2015) Inhibition of acid sphingomyelinase depletes cellular phosphatidylserine and mislocalizes K-Ras from the plasma membrane. Mol Cell Biol. 36, MCB.00719-15

11. Dharini van der Hoeven, Cho, K., Zhou, Y., Ma, X., Chen, W., Naji, A., Montufar-Solis, D., Zuo, Y., Kovar, S. E., Levental, K. R., Frost, J. A., van der Hoeven, R., and Hancock, J. F. (2017) Sphingomyelin Metabolism Is a Regulator of K-Ras Function. Mol Cell Biol. 10.1128/mcb.00373-17

12. del Vecchio, K., Frick, C. T., Gc, J. B., Oda, S. I., Gerstman, B. S., Saphire, E. O., Chapagain, P. P., and Stahelin, R. v. (2018) A cationic, C-terminal patch and structural rearrangements in Ebola virus matrix VP40 protein control its interactions with phosphatidylserine. Journal of Biological Chemistry. 293, 3335–3349

13. Adu-Gyamfi, E., Johnson, K. A., Fraser, M. E., Scott, J. L., Soni, S. P., Jones, K. R., Digman, M. A., Gratton, E., Tessier, C. R., and Stahelin, R. V. (2015) Host Cell Plasma Membrane Phosphatidylserine Regulates the Assembly and Budding of Ebola Virus. J Virol. 89, 9440–9453

14. Liu, X., Pappas, E. J., Husby, M. L., Motsa, B. B., Stahelin, R. v., and Pienaar, E. (2022) Mechanisms of phosphatidylserine influence on viral production: A computational model of Ebola virus matrix protein assembly. Journal of Biological Chemistry. 298, 102025

15. Husby, M. L., Amiar, S., Prugar, L. I., David, E. A., Plescia, C. B., Huie, K. E., Brannan, J. M., Dye, J. M., Pienaar, E., and Stahelin, R. V (2022) Phosphatidylserine clustering by the Ebola virus matrix protein is a critical step in viral budding. EMBO Rep. 10.15252/embr.202051709

16. Adu-Gyamfi, E., Digman, M. A., Gratton, E., and Stahelin, R. V. (2012) Investigation of Ebola VP40 assembly and oligomerization in live cells using number and brightness analysis. Biophys J. 102, 2517–2525

17. Adu-Gyamfi, E., Soni, S. P., Xue, Y., Digman, M. A., Gratton, E., and Stahelin, R. V. (2013) The ebola virus matrix protein penetrates into the plasma membrane: A key step in viral protein 40 (VP40) oligomerization and viral egress. Journal of Biological Chemistry. 288, 5779–5789

18. Hoenen, T., Biedenkopf, N., Zielecki, F., Jung, S., Groseth, A., Feldmann, H., and Becker, S. (2010) Oligomerization of Ebola Virus VP40 Is Essential for Particle Morphogenesis and Regulation of Viral Transcription. J Virol. 84, 7053–7063

19. Wan, W., Clarke, M., Norris, M. J., Kolesnikova, L., Koehler, A., Bornholdt, Z. A., Becker, S., Saphire, E. O., and Briggs, J. A. (2020) Ebola and Marburg virus matrix layers are locally ordered assemblies of VP40 dimers. Elife. 10.7554/eLife.59225

20. Zhao, D., and Moore, J. S. (2003) Nucleation–elongation: a mechanism for cooperative supramolecular polymerization. Org Biomol Chem. 1, 3471–3491

21. Doshi, U. R., and Muñoz, V. (2004) The principles of α-helix formation: Explaining complex kinetics with nucleation-elongation theory. Journal of Physical Chemistry B. 108, 8497–8506

22. Zhao, D., and Moore, J. S. (2003) Nucleation-Elongation Polymerization under Imbalanced Stoichiometry. J Am Chem Soc. 125, 16294–16299

23. Chatani, E., and Yamamoto, N. (2018) Recent progress on understanding the mechanisms of amyloid nucleation. Biophys Rev. 10, 527–534

24. Job, D., Valiron, O., and Oakley, B. (2003) Microtubule nucleation. Curr Opin Cell Biol. 15, 111–117

25. Cross, R. A., Geeves, M. A., and Kendrick-Jones, J. (1991) A nucleation-elongation mechanism for the self-assembly of side polar sheets of smooth muscle myosin. EMBO Journal. 10, 747–756

26. Hariadi, R. F. (2011) Non-equilibrium dynamics of DNA nanotubes, California Institute of Technology

27. Kirschner, D., Pienaar, E., Marino, S., and Linderman, J. J. (2017) A review of computational and mathematical modeling contributions to our understanding of Mycobacterium tuberculosis within-host infection and treatment. Curr Opin Syst Biol. 3, 170–185

28. Chang, S. T., Linderman, J. J., and Kirschner, D. E. (2005) Multiple mechanisms allow Mycobacterium tuberculosis to continuously inhibit MHC class II-mediated antigen presentation by macrophages. PNAS

29. Viceconti, M., Henney, A., and Morley-Fletcher, E. (2016) In silico clinical trials: how computer simulation will transform the biomedical industry. Int J Clin Trials. 3, 37

30. Zhang, J., and Muthukumar, M. (2009) Simulations of nucleation and elongation of amyloid fibrils. Journal of Chemical Physics. 10.1063/1.3050295

31. Soni, S. P., Adu-Gyamfi, E., Yong, S. S., Jee, C. S., and Stahelin, R. V. (2013) The Ebola virus matrix protein deeply penetrates the plasma membrane: An important step in viral egress. Biophys J. 104, 1940–1949

32. Schulz, M., Iwersen-Bergmann, S., Andresen, H., and Schmoldt, A. (2012) Therapeutic and toxic blood concentrations of nearly 1,000 drugs and other xenobiotics. Crit Care. 10.1186/cc11441

33. Lange, G., Mandelkow, E.-M, Jagla, A., and Mandelkow, E. (1988) Tubulin oligomers and microtubule oscillations: Antagonistic role of microtubule stabilizers and destabilizers. Eur J Biochem. 178, 61–69

34. Gc, J. B., Pokhrel, R., Bhattarai, N., Johnson, K. A., Gerstman, B. S., Stahelin, R. v., and Chapagain, P. P. (2017) Graphene-VP40 interactions and potential disruption of the Ebola virus matrix filaments. 493, 176–181

35. Ruedas, J. B., Ladner, J. T., Ettinger, C. R., Gummuluru, S., Palacios, G., and Connor, J. H. (2017) Spontaneous mutation at amino acid 544 of the Ebola virus glycoprotein potentiates virus entry and selection in tissue culture. J. Virol. 10.1128/JVI.00392-17

36. Maekawa, M., Lee, M., Wei, K., Ridgway, N. D., and Fairn, G. D. (2016) Staurosporines decrease ORMDL proteins and enhance sphingomyelin synthesis resulting in depletion of plasmalemmal phosphatidylserine. Sci Rep. 6, 1–14

37. Cho, K. J., Park, J. H., Piggott, A. M., Salim, A. A., Gorfe, A. A., Parton, R. G., Capon, R. J., Lacey, E., and Hancock, J. F. (2012) Staurosporines disrupt phosphatidylserine trafficking and mislocalize ras proteins. Journal of Biological Chemistry. 287, 43573–43584

38. Bennett, R. P., Finch, C. L., Postnikova, E. N., Stewart, R. A., Cai, Y., Yu, S., Liang, J., Dyall, J., Salter, J. D., Smith, H. C., and Kuhn, J. H. (2021) A novel ebola virus vp40 matrix protein-based screening for identification of novel candidate medical countermeasures. Viruses. 10.3390/v13010052

39. Kuge, O., Nishijima, M., and Akamatsu, Y. (1986) Phosphatidylserine Biosynthesis in Cultured Chinese Hamster Ovary Cells I. INHIBITION OF DE NOVO PHOSPHATIDYLSERINE BIOSYNTHESIS BY EXOGENOUS PHOSPHATIDYLSERINE AND ITS EFFICIENT INCORPORATION. Journal of Biological Chemistry. 261, 5784–5789

40. Vance, J. E., and Tasseva, G. (2013) Formation and function of phosphatidylserine and phosphatidylethanolamine in mammalian cells. Biochim Biophys Acta Mol Cell Biol Lipids. 1831, 543–554

41. Hasegawa, K., Kuge, O., Nishijima, M., and Akamatsu, Y. (1989) Isolation and characterization of a Chines hamster ovary cell mutant with altered regulation of phosphatidylserine biosynthesis. Journal of Biological Chemistry. 264, 19887–19892

42. Kuge, O., Saito, K., and Nishijima, M. (1999) Control of phosphatidylserine synthase II activity in Chinese hamster ovary cells. Journal of Biological Chemistry. 274, 23844–23849

